# Measurement of Atom Resolvability in CryoEM Maps with Q-scores

**DOI:** 10.1101/722991

**Authors:** Grigore Pintilie, Kaiming Zhang, Zhaoming Su, Shanshan Li, Michael F. Schmid, Wah Chiu

## Abstract

CryoEM density maps are now at the point where resolvability of individual atoms can be achieved. However, resolvability is not necessarily uniform throughout the map. We introduce a quantitative parameter to characterize the resolvability of individual atoms in cryoEM maps, the map Q-score. Q-scores can be calculated for atoms in proteins, nucleic acids, water, ligands, and other solvent atoms, using models fitted to or derived from cryoEM maps. Q-scores can also be averaged to represent larger features such as entire residues and nucleotides. Averaged over entire models, Q-scores correlate very well with the estimated resolution of cryoEM maps for both protein and RNA. Assuming the models they are calculated from are well-fitted to the map, Q-scores can thus be used as another measure to indicate resolvability of features in cryoEM maps at various scales, from entire complexes down to individual atoms. Q-score analysis of multiple cryoEM maps of the same proteins derived from different labs confirms reproducibility of structural features down to water and ion atoms.

## Introduction

CryoEM single particle methods strive to create accurate, high-resolution 3D maps of macromolecular complexes. Depending on many factors including imaging apparatus, detector, reconstruction method, structure flexibility, sample heterogeneity, and differential radiation damage, resulting maps have varying degrees of resolvability, or the level at which molecular features can be seen. Accurate quantification of resolvability in cryoEM maps has been a challenge in the field^1^. This task is very important as it can affect the interpretability and insights derived from such maps.

For every cryoEM map, a resolution is commonly reported or estimated, calculated from a Fourier shell correlation (FSC) plot between two independent reconstructions of the same complex^2^. It is well recognized that cryoEM maps usually do not have isotropic resolution throughout, hence a single number may not accurately represent the entire map. Local resolution can be estimated by most of image processing software (e.g. ResMap^3^), however such information is not as easy to comprehend in terms of specific residues as in the case with an atomic model.

Atomic models can be either fitted or built directly into cryoEM maps^4,5^. Map-model scores are then calculated from the model and map to assess how well the model fits the map^6^. Refinement^7^ or flexible fitting^8,9^ can then be applied, while making sure not to distort or overfit to noise^10,11^. The latter is accomplished by applying various stereochemical constraints, e.g. proper bond lengths, angles, dihedrals, preferred rotamers and van-der Waals distances; additional secondary-structure constraints (e.g.in the form of hydrogen bonds) can also be applied^7,9,12,13^.

Once an atomic model has been fitted to or derived from a cryoEM map, it can then be used to measure the resolvability of the features in the map. This can be done in several ways, including a map-model FSC curve, which requires that the model first be converted to a cryoEM-like map at the same resolution as the map. Occupancies and atomic displacement parameters of residues or atoms can also be used in this process to make the model-map better match the cryoEM map^14^. However, the FSC plot reflects the entire map volume. Proper masking may evaluate the resolvability of smaller features such as individual protein chains^10^, however it is impractical to quantify the resolvability of even smaller features such as a single side chain using this approach.

Two other methods that measure resolvability of such smaller features in a cryoEM map using a fitted model are EMRinger^15^ and Z-scores^16^. EMRinger considers map values near carbon-β atoms, while Z-scores can be applied to secondary structures or entire side chains. These scores were shown to correlate with the reported resolution when averaged over an entire map and model, meaning they can also be used to support the estimated resolution of the map. Moreover, they can also pinpoint smaller features in the model (e.g. secondary structures or side chains) which are not well-resolved in the map or not fitted properly to the map.

CryoEM maps have reached resolutions nearer to atomic-scale, for example apoferritin at 1.54Å (EMD:9865), 1.62Å (EMD:0144)^17^, 1.65Å (EMD:9599), and 1.75Å (EMD: 20026). A new question now arises as to how resolvability of individual atoms may be assessed. In crystallography, this is often reflected in the B-factor calculated for each atom^18^. Several formulations and interpretations of the B-factors are possible^19^, and their use in cryoEM has been suggested in the form of atomic displacement parameters (ADPs)^14^. So far, such formulations have not been fully characterized in terms of resolvability and resolution of map.

In this paper, we introduce a new score which is calculated from map values around an atom’s position, the Q-score. It aims to be a direct measure of the resolvability of atoms in cryoEM maps of complexes containing proteins, nucleic acids, and solvent molecules.

## Atomic Map Profiles

The basis of the Q-score is the atomic map profile. Atomic map profiles are calculated by averaging map values at increasing radial distances from the atom’s position. The radial distances range from 0Å to 2.0Å, and only points that are closer to the atom in question than to any other atoms in the model are considered. Figure 1A shows example atomic profiles in our two new maps of Apoferritin with resolutions of 1.75Å and 2.32Å, now deposited as EMD:20026, and EMD:20027. The model is the X-ray model of Apoferritin, (PDB:3ajo), which was first rigidly fitted to the cryoEM map, and then further refined using the Phenix real-space refinement procedure^7^. In the examples, atomic profiles have Gaussian-like contours. We consider a Gaussian equation of the form:

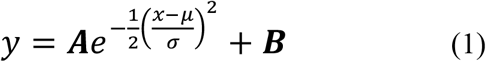

**Figure 1.**
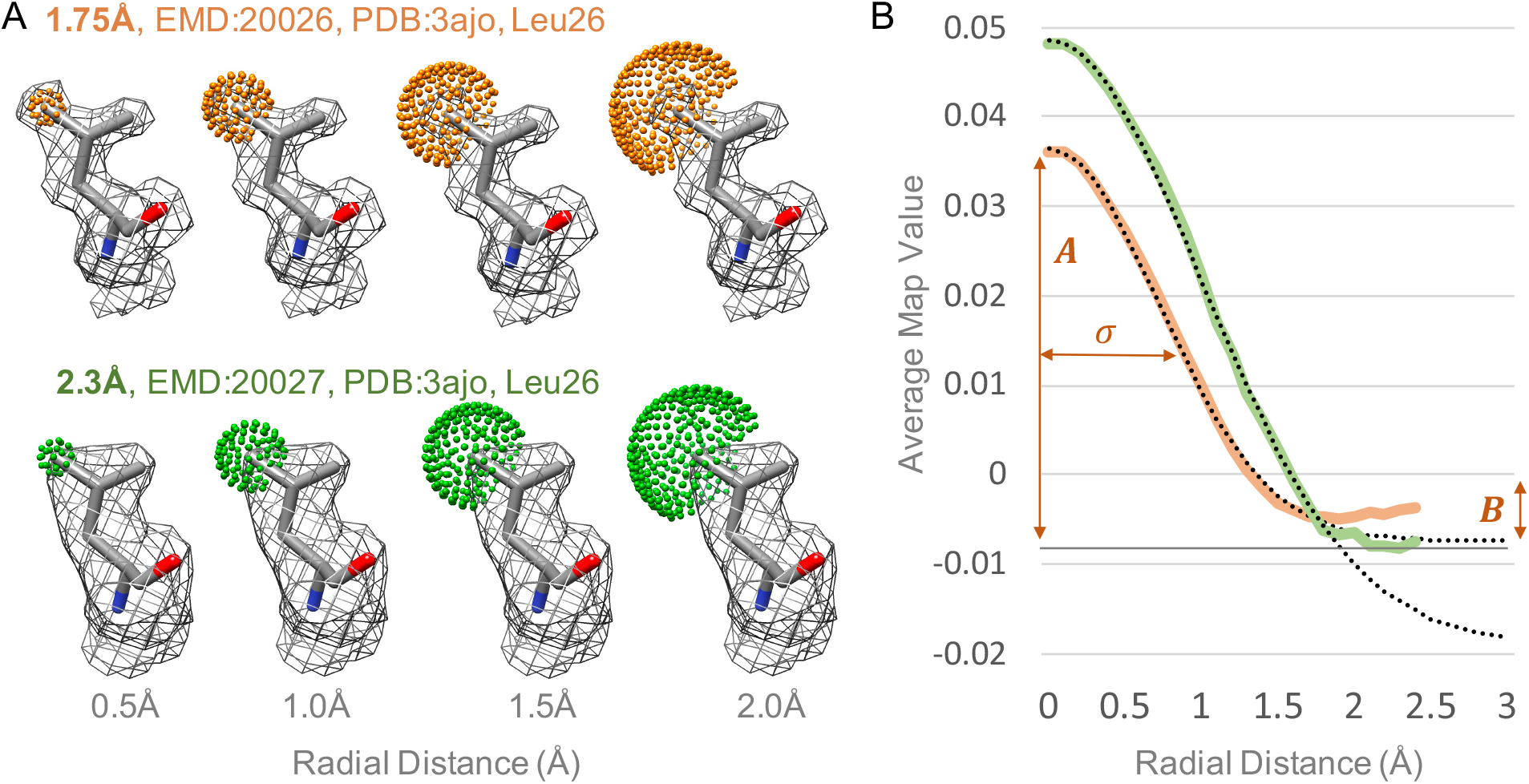
Atomic map profiles in cryoEM maps of Apoferritin at 1.75Å and 2.3Å resolution. (A) The residue Leu26 in the fitted model (PDB:3ajo) is shown, along with contour surface of the cryoEM map around this residue. Spherical shells of points centered on the CD2 atom are shown at increasing radial distances; only points that are closer to the CD2 atom than to any other atom in the model are used. (B) Average map values at these points are plotted vs. radial distance; these are the atomic map profiles. The dotted lines represent Gaussian functions with parameters ***A***, ***B*** and ***σ*** which are fitted to each profile.

Gaussian functions of the form in Eqn.1, where *x* is the radial distance and *y* the average map value, fit extremely well to the atomic profiles shown in Figure 1, up to a distance of 2Å (mean error of ~0.01Å). Past this distance, observations in various maps indicate that atomic map profiles become noisy and start to increase. This is likely due to effects from other nearby atoms and/or solvent.

When the model is well-fitted to the map, the relative height, ***A-B***, and width, ***σ***, of the Gaussian function (Eqn.1) fitted to the profile may be considered to be proportional to several factors including the resolution of the map, and the overall mobility of the atom. It may be impossible to fully separate such factors based on the observed cryoEM map alone. Regardless of the cause, the overall Gaussian profile seen in the map represents to what degree the respective atom is resolved in the map - the more resolved an atom is in the map, the higher (relative to other peaks in the same map) and narrower (up to a certain point, i.e. the radius of the atom itself) the Gaussian profile around it would be.

## Q-score

The idea behind the Q-score is to measure how closely the map profile of an atom matches that of the Gaussian-like function we would see if an atom is well-resolved. Thus, to calculate the Q-score, the atomic map profile is compared to a ‘reference Gaussian’ as given by Eqn. 1, with the following parameters:

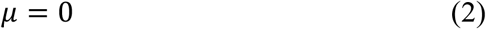

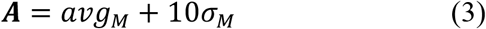

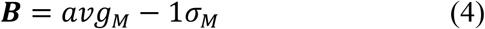

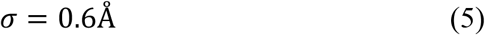

In the above, the mean, μ, is set to 0, as the Gaussian is expected to be centered around the atom’s position. The parameters ***A*** and ***B*** are obtained using the mean/average across all values in the entire map, *avg*_*M*_, and the standard deviation of all values around this mean, *σ*_*M*_. A well resolved atom would be centered on a peak that has a relatively high value in the map, and fall off to a value below the mean, but not necessarily as low as the background noise. The width of the reference gaussian is set as σ=0.6. These parameters in Eqns. 2–5 are chosen to make the reference Gaussian roughly match the atomic profile of a well-resolved atom in the 1.54Å cryoEM map as shown in Figure 2B The height of the reference Gaussian is different in each map, accounting for differences in the range of map values often seen in different maps; for example, in Figure 1B, the values in in 2.3Å map are higher than those in the 1.75Å map.

**Figure 2.**
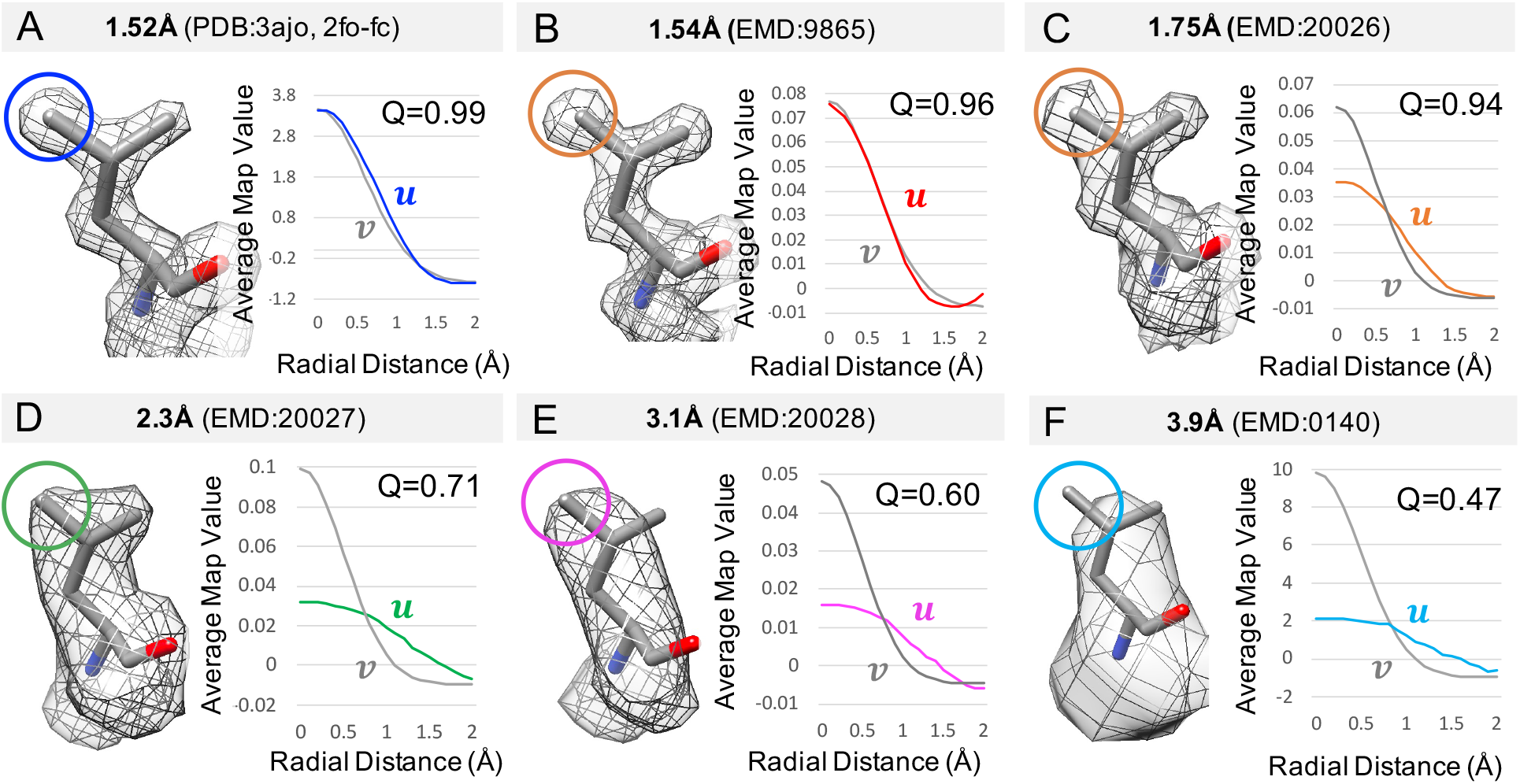
Calculation of Q-scores for an atom in 6 maps at different resolutions, including an X-ray map. The atom is CD2 from Leu 26 in PDB:3ajo. The atomic profile in each map is marked with the letter ***u***, while the reference Gaussian is marked with ***v***.

The Q-score is then calculated as a correlation between values in the atomic profile obtained from the map, ***u***, and values obtained from the reference Gaussian, ***v***, defined in Eqn. 1 and with parameters in Eqns. 2–5. The following normalized, about the mean, cross-correlation formula is used:

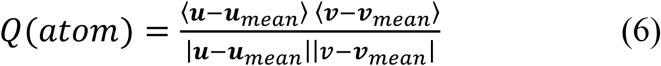

Several atomic profiles and reference Gaussians are illustrated in Figure 2, for an X-ray map and 5 cryoEM maps at various resolution. At high resolutions, the atomic profiles are more similar to the reference Gaussian, and hence Q-scores are higher. At lower resolutions, the atomic profile of the same atom is wider than the reference Gaussian, hence Q-scores are lower. Q-scores would also be low for atomic profiles that are mostly noise (e.g. random values or a sharp peak). In some cases when the atom is not well-placed in the map, the Q-score can be negative if the atomic profile has a shape that increases away from the atom’s position.

Calculating Q-scores is similar to calculating a cross-correlation between the model and a cryoEM map, using a simulated map of the model blurred using a Gaussian function with the parameters in Eqns. 2–5. The main difference is that with Q-scores, the cross-correlation is performed atom-by-atom, separating out parts of the density that are closest to each atom. The cross-correlation about the mean is used so that the Q-scores decrease as resolution also decreases. When not subtracting the mean, this effect would not be ensured^16^.

## Q-scores of Atoms in Proteins

Figure 3 shows Q-scores for atoms taken from maps of Apoferritin at various resolutions. One of the maps is an X-ray map at 1.52Å resolution (2fo-fc, PDB:3ajo) as a reference; another is a recent high-resolution map at 1.54Å (EMD:9599). The other three are new maps we reconstructed to 1.75Å (EMD:20026), 2.3Å (EMD:20027), and 3.1Å (EMD:20028) with different numbers of particle images, from the same data set. For the cryoEM maps, the X-ray model PDB:3ajo was fitted to the density and also refined using Phenix real-space refinement^7^. Q-scores for each atom correlate well with visual resolvability, i.e. the more resolvable an atom, the higher the Q-score. They also increase as the estimated resolution of the map increases.

**Figure 3.**
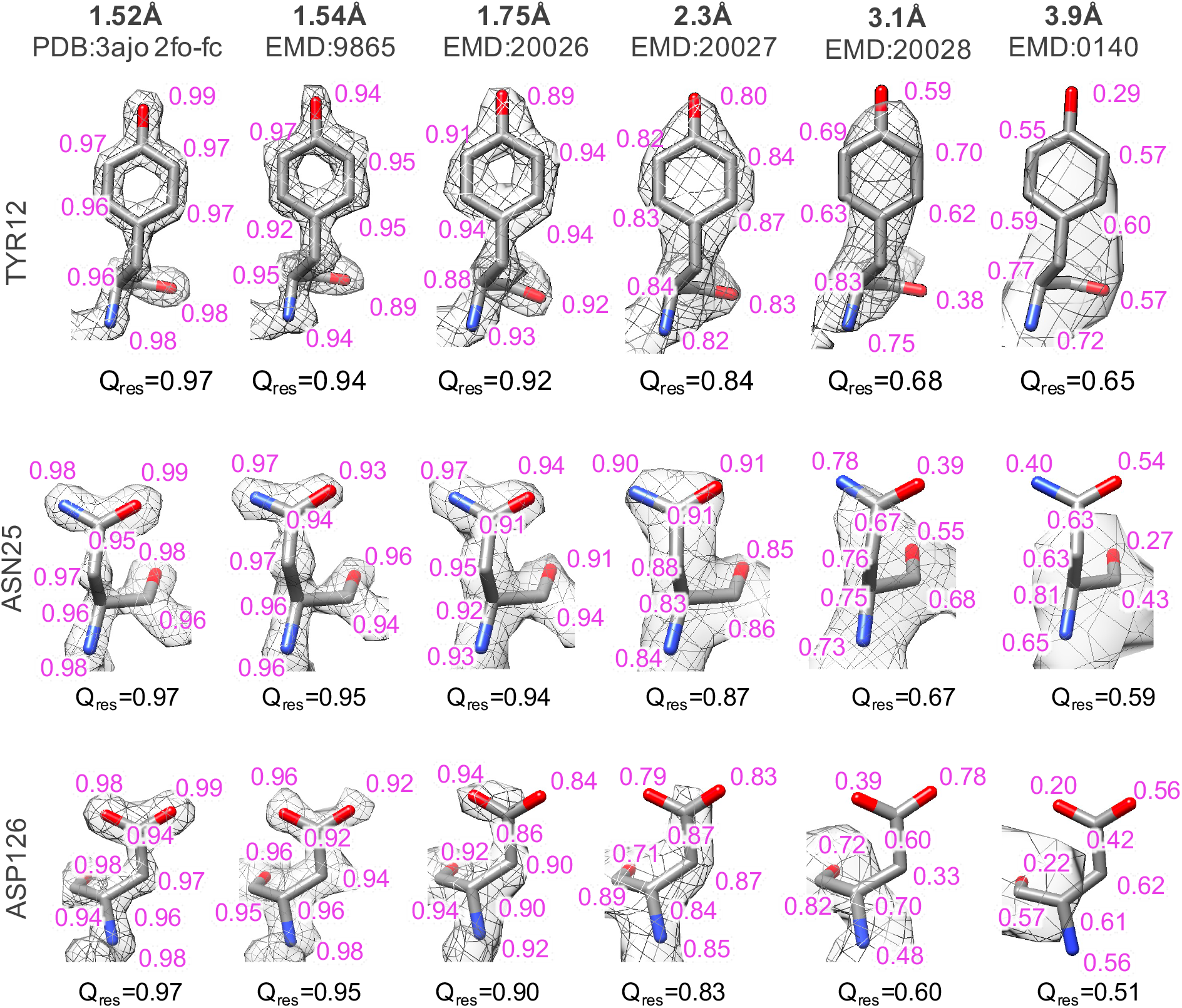
Atom Q-scores for three types of residues, taken from Apoferritin maps at various resolutions. Atom Q-scores are shown in purple close to each atom.

Resolvability and Q-scores can decrease for some residues faster than others as a function of resolution. For example, in Figure 3, the Q-score for ASP126 drops more than for ASN25 from 1.52Å to 3.9Å. This effect may be due to several reasons. First, some residue types may be more susceptible to radiation damage (as previously shown using EMRinger^15^). Also, certain residue types may be more conformationally dynamic, or occur in environments that are more dynamic (e.g. solvent accessible), and hence may not resolve as well with a fewer number of particles. Finally, the interaction of the electron beam with charged side chains may have a weakening effect on map values around them^14^.

## Q-scores for Atoms in Nucleic Acids

Q-scores can also be calculated for atoms in models of nucleic acids. In figure 4, we used several maps and models containing RNA from the EMDB at resolutions ranging from 2.5Å to 4.0Å. Q-scores were averaged over atoms in bases, phosphate-sugar backbones, and entire nucleotides. As with proteins, Q-scores decrease with resolvability and estimated map resolution.

**Figure 4.**
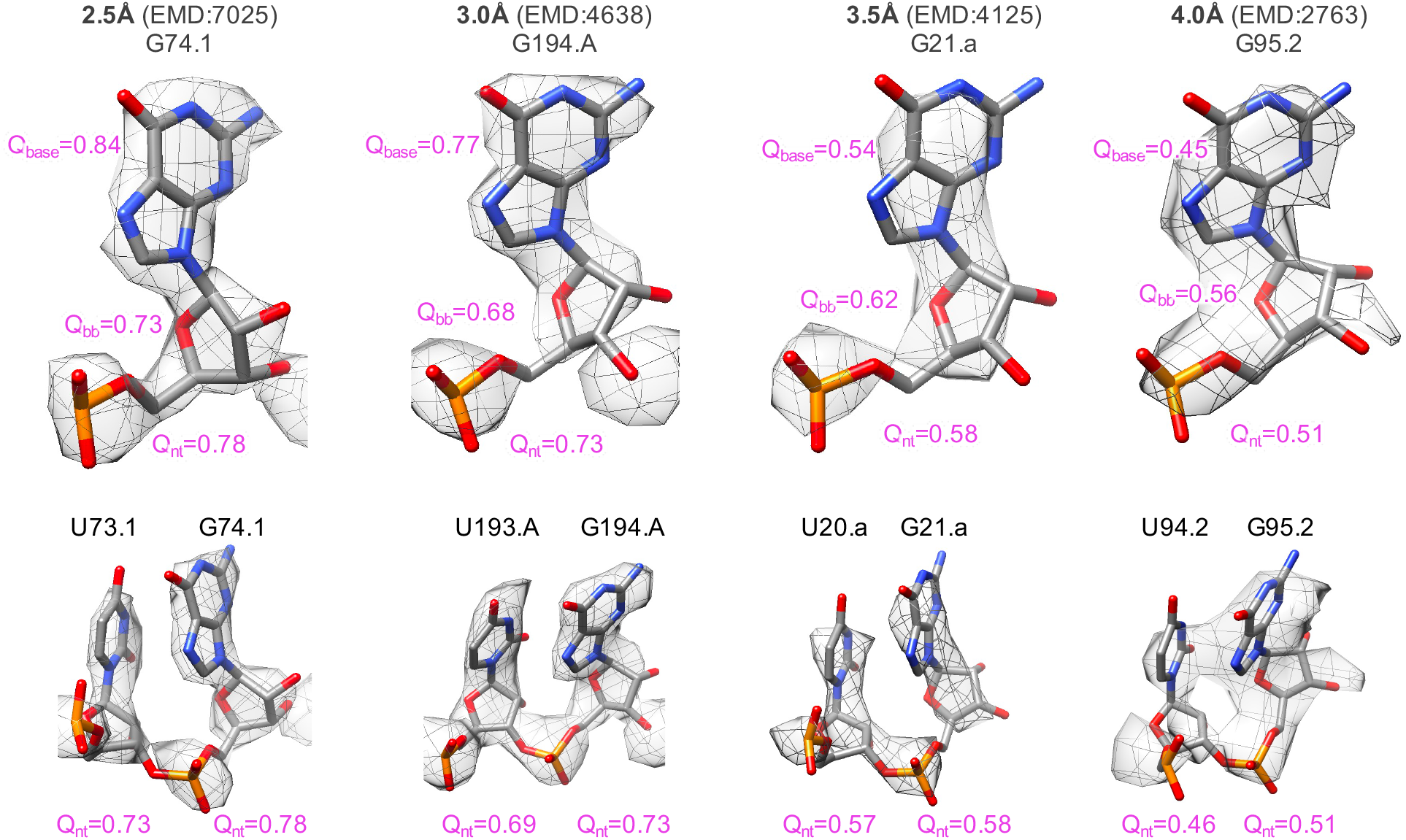
Q-scores averaged over entire nucleotides (Q_nt_) in RNA maps and models from the EMDB at four different resolutions. Q-scores are also averaged for the base (Q_base_) and phosphate-sugar backbone (Q_bb_) groups in the nucleotides shown on the top row.

Figure 4 also illustrates a general trend that at ~4Å and lower resolutions, stacked bases from adjacent nucleotides are typically not separable in cryoEM maps, whereas at higher than 4Å resolutions, they usually do become separate at appropriate contour levels.

It is also interesting to note that for the examples in Figure 4, at high resolutions (~2.5Å), the difference in Q-score or resolvability of individual bases is higher than that of the backbone (0.84 for base vs. 0.73 for backbone). Going towards lower resolutions in this example, bases become less resolvable (0.45 for bases vs 0.56 for backbone). This may be counter-intuitive as bases can have higher values in the map (i.e. appear first at a high contour level). However, these contours may have overall less detail as adjacent stacked bases are not fully separable and merge together.

## Q-score vs. Resolution

Q-scores can also be averaged across an entire model to represent an average resolvability measure for the entire map. Such average Q-scores were plotted as a function of reported resolution for a number of maps and models obtained from the EMDB. Figure 5 shows these plots for two sets of maps and models, one set using only protein models, and the other set only nucleic acids (RNA). The protein set includes the maps used in the EMRinger analysis^15^, and further adding 24 maps of Apoferritin and β-galactosidase at resolutions up to 1.54Å. In the RNA set, a total of 52 maps and models were used at a range of resolutions ranging from 2.5Å (the highest resolution of an RNA-containing map to date) to 5.4Å. The full sets are listed in Tables 1 and 2. In both cases, the average Q-score correlates very strongly to reported resolution, with R^2^ of 0.90 for proteins and 0.89 for RNA. The R^2^ quantifies the error in fitting the linear function to the observed data; it is 1 for perfect correlation and 0 for no correlation. The high values of R^2^ in Figure 5 show that Q-scores closely capture the resolvability of atomic features in cryoEM maps. Thus, average Q-scores from a properly fitted model may be useful as a measure of resolvability in the map in addition to the reported resolution.

**Figure 5.**
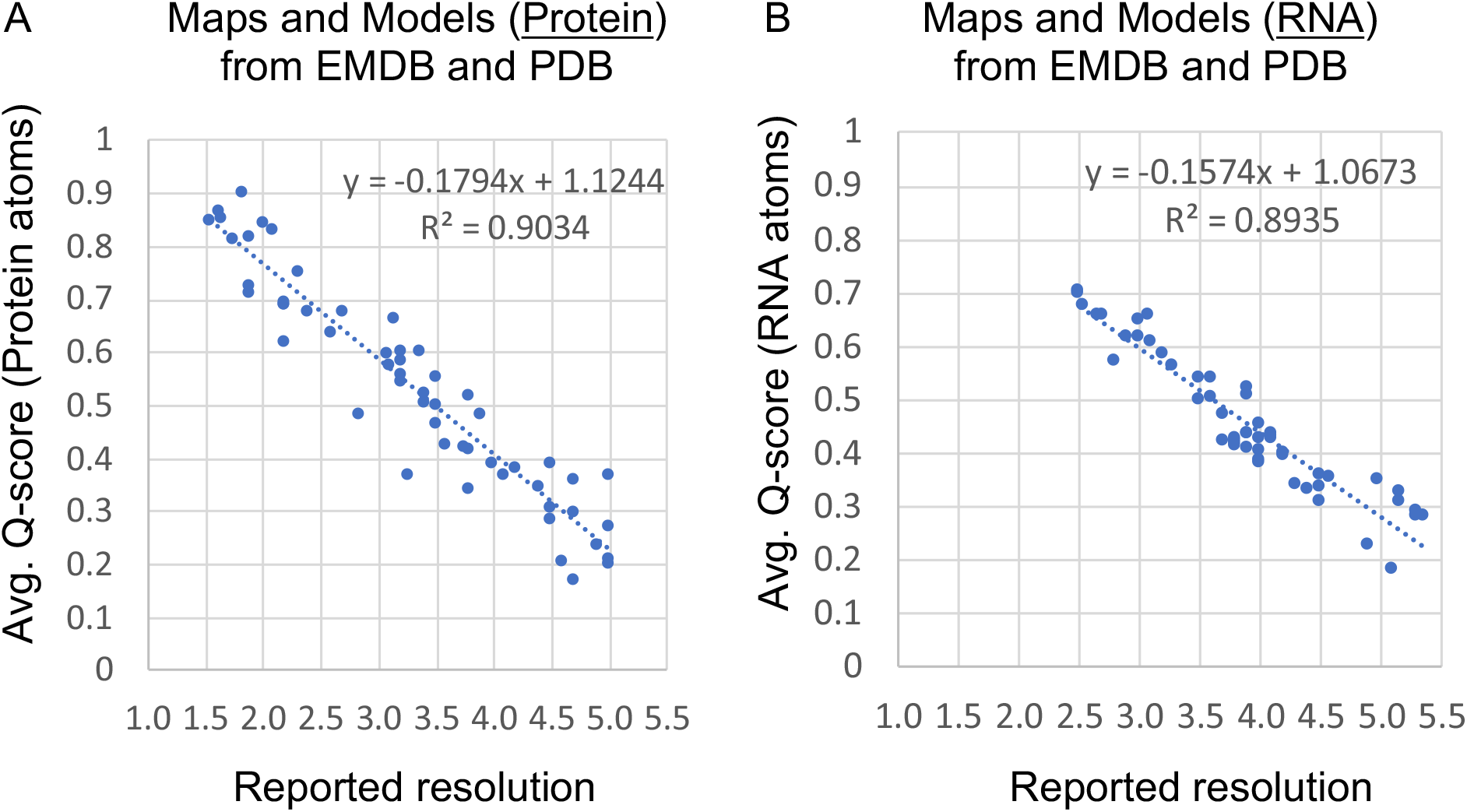
Model Q-scores compared to reported resolution for maps and models obtained from EMDB. (A) Average Q-scores for atoms in proteins. (B) Average Q-scores for atoms in RNA.

**Table 1.**
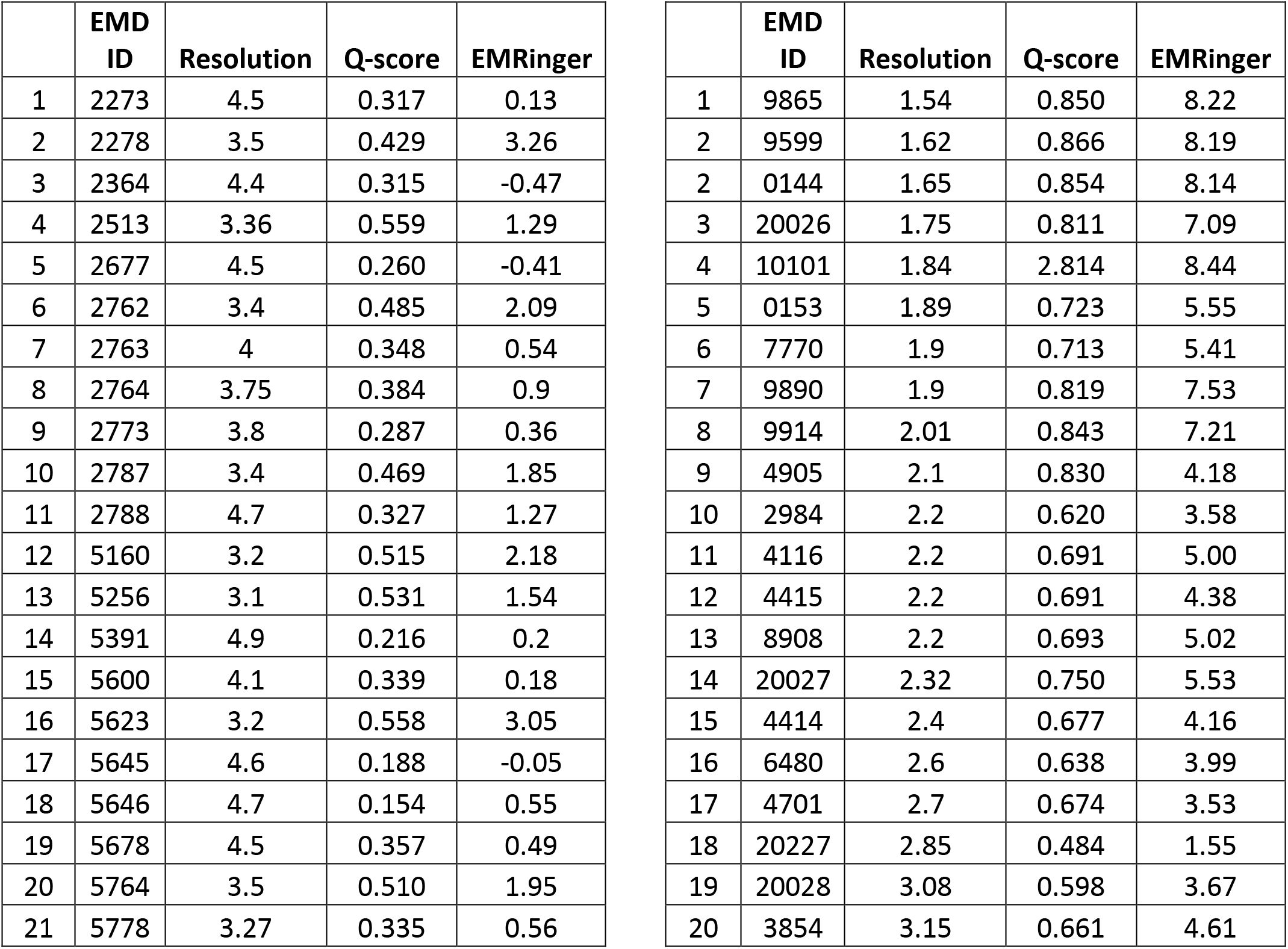
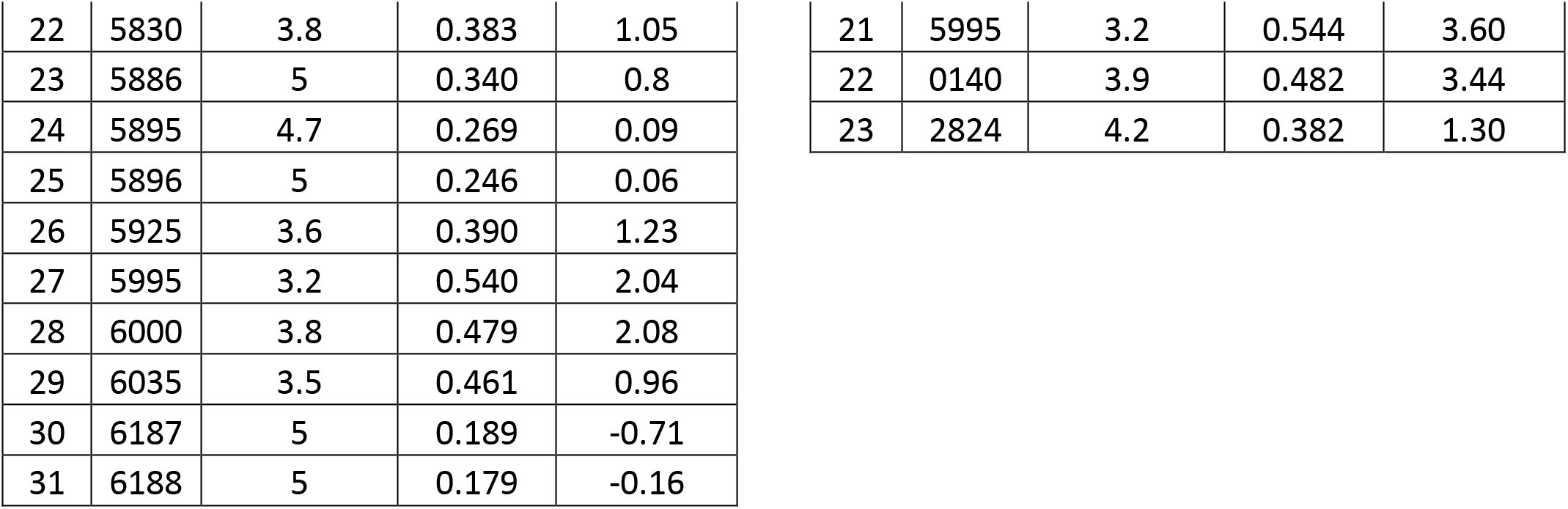
Maps from EMDB for which Q-scores of protein components are calculated for the plot in Figure 5A. The table on the left shows the original maps used in the original EMRinger analysis^15^. The table on the right contains a new set of maps of Apoferritin and β-galactosidase.

**Table 2.** Maps from EMDB containing RNA for which Q-scores vs. resolution are plotted in Figure 5B.

|  | EMD ID | PDB File | Resolution | Q-score |
| --- | --- | --- | --- | --- |
| 1 | 7025 | 6az3-pdb-bundle2 | 2.5 | 0.699649 |
| 2 | 7025 | 6az3-pdb-bundle1 | 2.5 | 0.703164 |
| 3 | 8361 | 5t5h-pdb-bundle1 | 2.54 | 0.675016 |
| 4 | 0243 | 6hma | 2.65 | 0.656016 |
| 5 | 7024 | 6az1 | 2.7 | 0.656345 |
| 6 | 6583 | 3jcs-pdb-bundle1 | 2.8 | 0.573334 |
| 7 | 20173 | 6ore-pdb-bundle1 | 2.9 | 0.615376 |
| 8 | 4638 | 6qul | 3 | 0.649598 |
| 9 | 0600 | 6ole-pdb-bundle3 | 3 | 0.618584 |
| 10 | 0233 | 6hiz-pdb-bundle1 | 3.08 | 0.655904 |
| 11 | 4560 | 6qik-pdb-bundle1 | 3.1 | 0.606439 |
| 1 | 10068 | 6rzz-pdb-bundle1 | 3.2 | 0.584933 |
| 2 | 0101 | 6gzq-pdb-bundle1 | 3.28 | 0.564434 |
| 3 | 4125 | 5lze-pdb-bundle1 | 3.5 | 0.497689 |
| 4 | 4125 | 5lze-pdb-bundle2 | 3.5 | 0.538745 |
| 5 | 2938 | 4ug0-pdb-bundle1 | 3.6 | 0.54117 |
| 6 | 2938 | 4ug0-pdb-bundle2 | 3.6 | 0.503634 |
| 7 | 6559 | 3jcj-pdb-bundle1 | 3.7 | 0.471255 |
| 8 | 6559 | 3jcj-pdb-bundle2 | 3.7 | 0.421646 |
| 9 | 8620 | 5uyq-pdb-bundle1 | 3.8 | 0.42095 |
| 10 | 8620 | 5uyq-pdb-bundle2 | 3.8 | 0.426834 |
| 11 | 0076 | 6gwt-pdb-bundle2 | 3.8 | 0.417034 |
| 12 | 0076 | 6gwt-pdb-bundle1 | 3.8 | 0.411493 |
| 13 | 0192 | 6hcf-pdb-bundle2 | 3.9 | 0.524457 |
| 14 | 0192 | 6hcf-pdb-bundle1 | 3.9 | 0.506735 |
| 15 | 0192 | 6hcf-pdb-bundle3 | 3.9 | 0.408919 |
| 16 | 8279 | 5kps-pdb-bundle2 | 3.9 | 0.43481 |
| 17 | 8279 | 5kps-pdb-bundle1 | 3.9 | 0.437756 |
| 18 | 8618 | 5uyn-pdb-bundle2 | 4 | 0.383391 |
| 19 | 8618 | 5uyn-pdb-bundle1 | 4 | 0.385951 |
| 20 | 4080 | 5lmu | 4 | 0.428174 |
| 21 | 2763 | 3j81_real_space_refined | 4 | 0.453 |
| 22 | 2763 | 3j81 | 4 | 0.402465 |
| 23 | 4350 | 6g51 | 4.1 | 0.426988 |
| 24 | 8280 | 5kpv-pdb-bundle1 | 4.1 | 0.435967 |
| 25 | 8280 | 5kpv-pdb-bundle2 | 4.1 | 0.43031 |
| 26 | 643 | 6o7k | 4.2 | 0.395112 |
| 27 | 20188 | 6ost-pdb-bundle1 | 4.2 | 0.3981 |
| 28 | 4382 | 6gc7 | 4.3 | 0.339055 |
| 29 | 0083 | 6gxp-pdb-bundle1 | 4.4 | 0.331033 |
| 30 | 4349 | 6g4w | 4.5 | 0.311581 |
| 31 | 3133 | 5ady | 4.5 | 0.361631 |
| 32 | 4351 | 6g53 | 4.5 | 0.338915 |
| 33 | 0104 | 6gzx-pdb-bundle1 | 4.57 | 0.356781 |
| 34 | 4083 | 5lmv | 4.9 | 0.228577 |
| 35 | 3553 | 5mrf-pdb-bundle1 | 4.97 | 0.349647 |
| 36 | 8473 | 5tzs | 5.1 | 0.183097 |
| 37 | 3661 | 5no2 | 5.16 | 0.326795 |
| 38 | 3662 | 5no3 | 5.16 | 0.310096 |
| 39 | 4122 | 5lzb-pdb-bundle1 | 5.3 | 0.28398 |
| 40 | 4427 | 6i7o-pdb-bundle1 | 5.3 | 0.292207 |
| 41 | 4075 | 5lmp | 5.35 | 0.282151 |

## Q-scores of Solvent Atoms

The X-ray Apoferritin model (PDB:3ajo) contains one protein chain, 229 oxygen (O) atoms (from water) and 12 Mg atoms. A closeup on the map and model with two Mg and three O atoms is shown in Figure 6. Q-scores calculated for each of these atoms correlate well with the contours seen in the map. Of the four maps shown, three of them are cryoEM maps at near-atomic resolutions (1.54Å, 1.65Å, and 1.75Å). The model used all cases comes from the X-ray map. It is reassuring to see that some of the solvent atoms placed in the X-ray map can also be observed in the cryoEM maps (e.g. Mg183, O280, O236). However, some of the solvent atoms (e.g. Mg184), is not seen equally well in all three maps; for example, in the 1.54Å and 1.65Å maps, Mg184 has low Q-score (0.12 and 0.03 respectively), and are not seen at the same map contour level where the other solvent atoms are seen.

**Figure 6.**
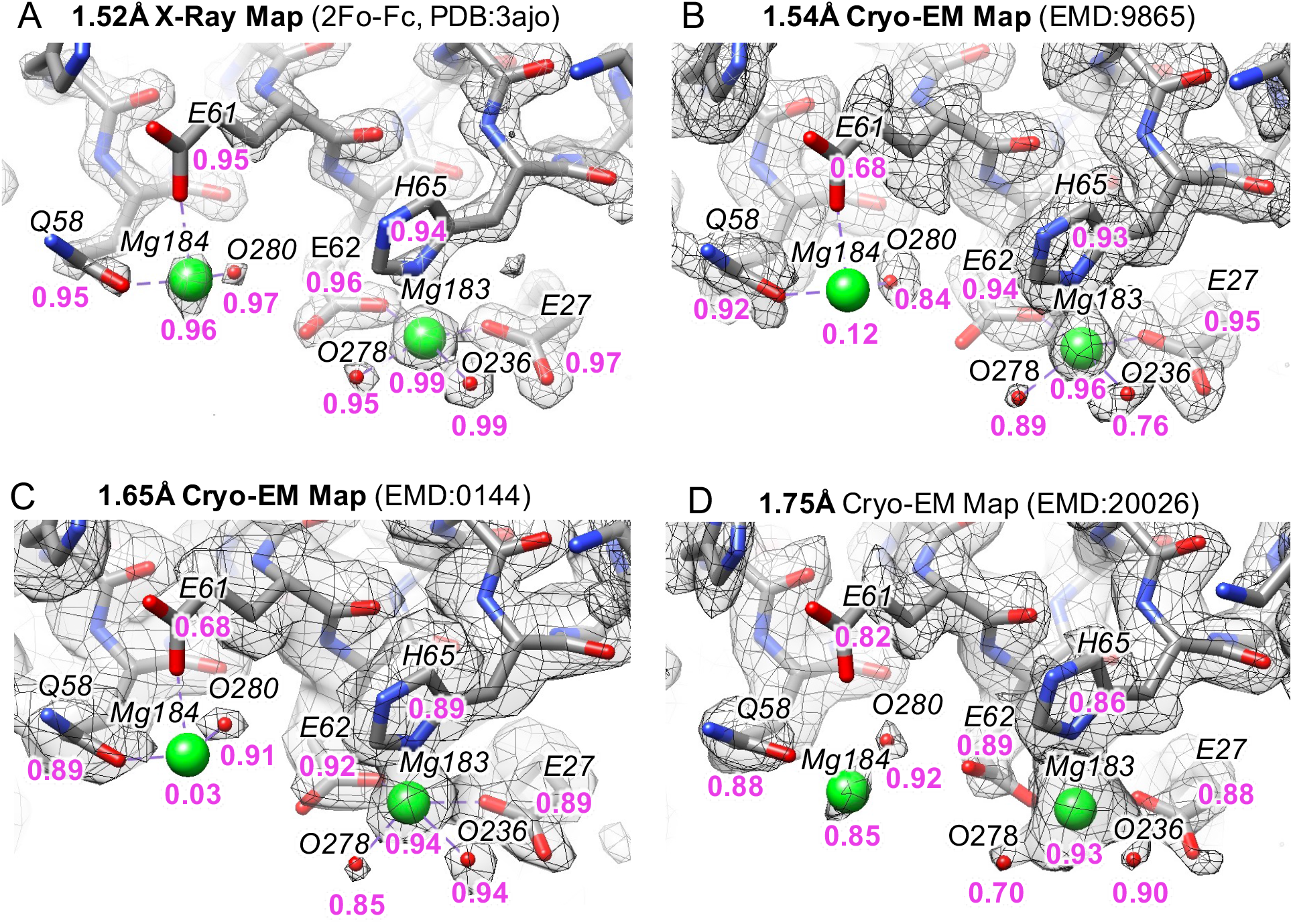
A close up in Apoferritin models showing solvent atoms (Mg and O from water), along with calculated Q-scores in purple under each atom and nearby residue. The model comes from the X-ray map (PDB:3ajo) shown in A. It was further refined into each of the three cryoEM maps, B-D.

In this region of the map, the three water molecules shown in Figure 6 have high Q scores and observable map contours. Along with Mg183, these provide evidence that cryoEM structures can be used to identify locations of solvent molecules, much like with X-ray crystallography. However, since Mg184 is only visible and has good Q-scores in only 2 of the 4 maps considered here, differences between the cryoEM and X-ray maps can also be seen. Such differences may be due to different affinities at some sites and/or different biochemical conditions across the different data sets.

Figure 7A shows distributions of Q-scores for solvent atoms in the X-ray map (PDB:3ajo). Most solvent atoms have very high Q-scores of 0.9 and higher. Visual inspection confirmed that all these solvent atoms can be seen in the X-ray map (2fo-fc), e.g. as shown in Figure 6A. Figure 7B,C shows distribution plots for the same model fitted to the cryoEM maps at 1.54Å and 1.75Å resolution, using the rigidly fitted model and also after refinement (including solvent atoms) of the rigidly fitted model using Phenix real-space refine^7^.

**Figure 7.**
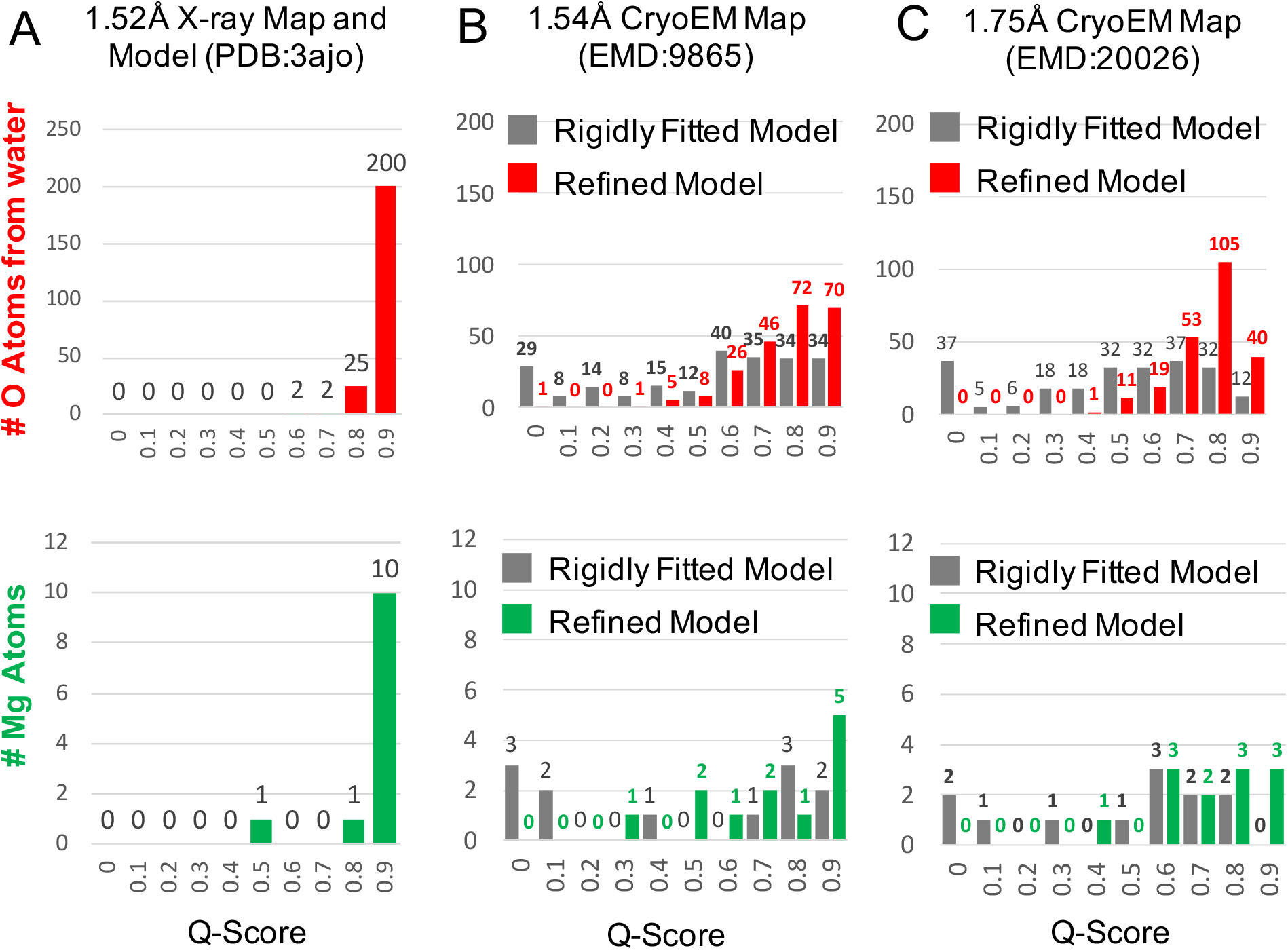
Distribution of Q-scores for solvent atoms (water and Mg) in X-ray map (PDB:3ajo), and in two cryoEM maps before and after refinement.

For the rigidly fitted model, Q-scores of the solvent atoms are considerably lower than in the X-ray map (Figure 7B). For example, in the 1.75Å cryoEM map, only 12 O atoms from water have Q-scores of 0.9 and higher, and 32 have Q-scores of 0.8 to 0.9. In the 1.54Å map, 34 atoms have Q-scores of 0.9 and higher, and another 34 have Q-scores of 0.8 to 0.9. Thus, water atoms are less resolved in the cryoEM maps than in X-ray. It is possible that some of the solvent atoms seen in the X-ray model may not be resolvable in the cryoEM maps or may be in different positions.

To explore whether solvent atoms may have different positions in the cryoEM maps, Q-scores of the solvent atoms were also calculated in the X-ray model after real-space refinement with Phenix^7^. This refinement method moves solvent atoms towards higher map values, while keeping them within reasonable distance of other atoms. The distributions in the Q-scores for solvent atoms after this procedure are plotted in Figure 7B, C for the two cryoEM maps. Q-scores are now higher; 142 water atoms in the 1.54Å map and 145 atoms in the 1.75Å map have Q-scores of 0.8 and higher, compared to 225 water atoms in the X-ray map with Q-scores of 0.8 and higher.

In the 1.54Å map, after refinement, water atoms with Q-scores 0.8 and higher moved between 0.1Å and 2.2Å, on average 0.54Å. In the 1.75Å map, the water atoms with Q-scores of 0.8 and higher moved between 0.1Å and 1.6Å, on average 0.67Å. Although it is difficult to assess the exact cause of the movements in these maps, it is reasonable to conclude that the water found in cryoEM maps are real and potentially within experimental errors of their atom positions in both X-ray and cryoEM structures.

In the above analysis, the water molecules were based on those originally observed in the X-ray map. If one studies a *de novo* map, the identification of water molecules would require a protocol used in modeling software, e.g. Phenix and Coot. In addition to such a protocol, Q-scores may be used as an additional validation parameter to assist in the finding of water and ions.

## Radial Plots for Solvent Atoms

Radial plots in Figure 8 further characterize distances between solvent atoms (H_2_O and Mg) and other atoms including other water molecules (H_2_O-H_2_O), and also O and N atoms in protein, H_2_O-O and H_2_O-N respectively. The radial plot for the X-ray model of Apoferritin (PDB:3ajo) is shown in Figure 8A. This plot shows that H_2_O-H_2_O distances have a sharp peak at 2.8Å. A similar peak is seen for distances between O atoms in water and O atoms in protein (H_2_O-O). Distances from Mg atoms to H_2_O and to O have smaller peaks (since there are much fewer Mg atoms in the model) at a distance of 2.2Å.

**Figure 8.**
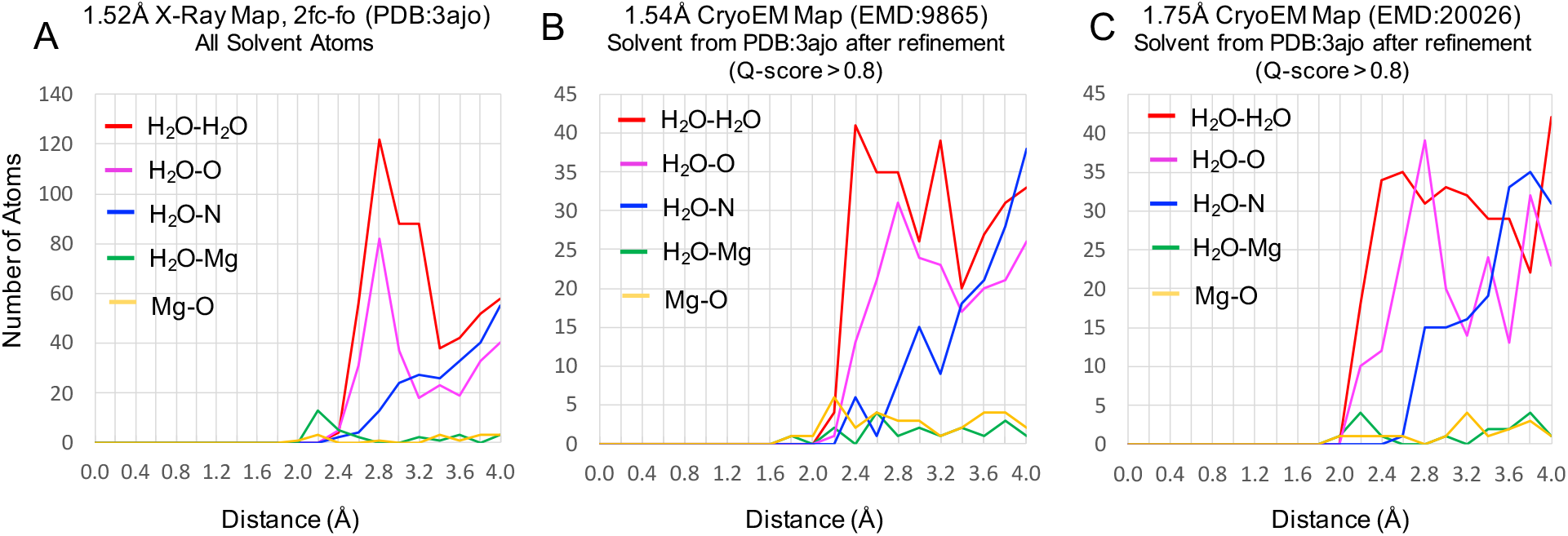
Radial plots of distances from solvent atoms to other types of atoms. Oxygen atoms in water are labeled H_2_O, whereas oxygen/nitrogen atoms in protein are labeled O/N respectively.

Radial plots for the X-ray model fitted to the 1.54Å and 1.75Å cryoEM maps are shown in Figure 8B, C, considering only solvent atoms with Q-scores of 0.8 and higher after refinement. A wider range H_2_O-H_2_O can be seen in both cases; instead of a sharp peak at 2.8Å, a broader peak from ~2.4Å up to ~3.2Å can be seen. This seems to indicate that water-water distances in Cryo-EM may vary. On the other hand, H_2_O-O still have a main peak at 2.8Å after refinement in both cryoEM maps, matching the peak seen in the X-ray map. Thus, H_2_O-O distances are very similar in X-ray and cryoEM maps in these examples, however the differences in H_2_O-H_2_O distances suggests that there may be a difference in water organization around protein in cryoEM vs X-ray maps.

## Q-scores of Solvent Atoms at different resolutions

Finally, we looked at the resolvability and Q-scores of solvent atoms in cryoEM maps of Apoferritin at different resolutions, as shown in Figure 9. The locations of the solvent atoms are again taken from the X-ray model (PDB:3ajo). As Figure 9 shows, Mg183 appears resolved at both 1.75Å and 2.3Å, with separable contours in both maps and high Q-scores (0.93 and 0.80). However, the contours no longer have a symmetric spherical shape, indicating possibly more variation in its position. In the 3.1Å map, the contour is no longer separable from that of the nearby His65 residue, and the Q-score is also considerably lower (0.60). The water atoms are similarly resolved in the 1.75Å and 2.3Å maps and contours around them can be seen, however at 3.1Å and 3.9Å they can no longer be seen and Q-scores become very low (−0.44 to 0.38).

**Figure 9.**
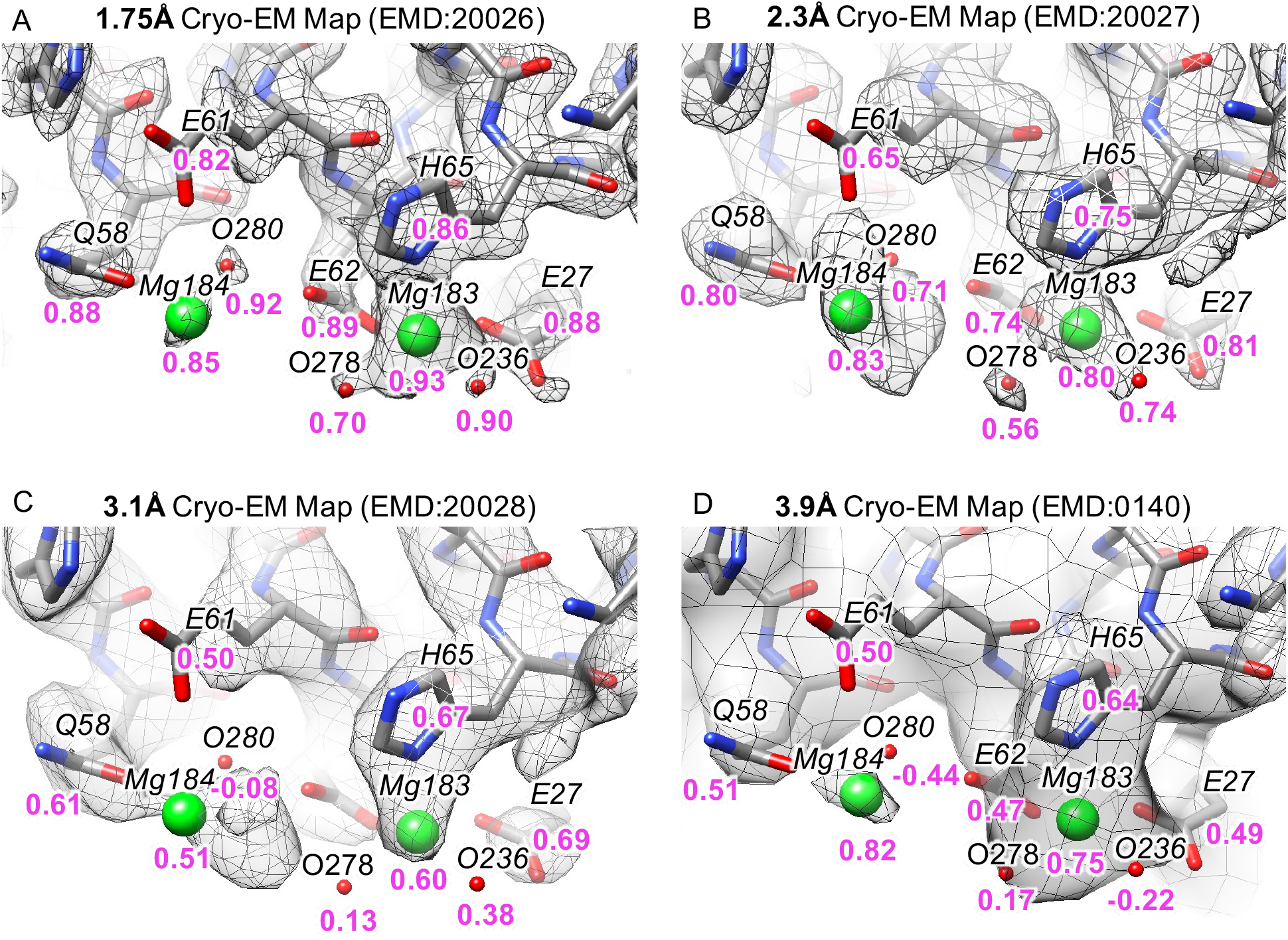
Solvent atoms from X-ray model (PDB:3ajo) in cryoEM maps at resolutions of 1.75Å to 3.9Å. Q-scores are shown in purple below each atom. Nearby residues with Q-scores are also labeled (Q58, E61, E62, H65, E27).

At 3.9Å resolution, both Mg atoms still have high Q-scores and thus high map values around them, and they can be seen at a lower threshold. However, the map contours at these thresholds do not necessarily separate them fully from the nearby residues (as again for Mg183). Nevertheless, even at such lower resolutions (3Å-4Å), it appears that the larger solvent atoms can still significantly influence the cryoEM map values, producing strong though more diffuse peaks in the map. This may have some implications when creating or refining models in such maps. Perhaps placement of solvent atoms should be considered for the model to be accurately created and/or refined.

## Conclusions

Q-scores can measure the resolvability of individual atoms in cryoEM maps, using atomic positions and nearby map values. As was noted, this metric is closely related to the map-model cross-correlation score, which is already widely used in the field to assess the fit of a model to a map. However, the Q-score improves in two ways on the cross-correlation score: 1) it is formulated so that it correlates to the resolution of the map and 2) it makes it applicable to small features (individual atoms) while avoiding explicit masking. Aside from this, it is important to note that nothing is assumed about the model itself, e.g. whether it has good stereochemistry; this could be deduced with other scores such as the Molprobity score^20^.

Q-scores averaged over entire models were shown to correlate very well with the reported resolution of cryoEM maps containing both proteins and nucleic acids. Various visualizations also showed that Q-scores indeed correlate well with the resolvability of individual atoms, and also groups of atoms such as side chains. However, it still requires a model to first be fitted to or built based on the cryoEM map. The score can be very useful to analyze the map and its resolvability in different regions, and also test whether the model accurately interprets the map. It could thus be useful as a map-model validation metric.

In this paper, several quantifiable observations were made with the help of Q-scores. For example, when applied to atoms in protein side chains, the Q-score showed that resolvability of certain types of side chains (Asp) drop faster than others (Asn) as a function of resolution. In the case of nucleic acids, per-nucleotide Q-scores could be used to indicate whether stacked bases are separable. Finally, Q-scores were also applied to water and other solvent atoms, helping to confirm that water and other solvent atoms can indeed be resolved and placed in cryoEM maps much as they are in X-ray crystallography.

## Experimental Methods

Data: all the data for the analysis were drawn from EMDB and PDB. The EMDB 20026, 20027, 20028 maps were collected in Titan Krios electron microscope (Thermo Fisher) at 300 keV equipped with BioQuantum energy filter and K2 director detector (Gatan). Images were recorded in movie mode and corrected prior to image processing with Relion software^17^. The map resolution was estimated from two independent maps with a total of 70,000 particle images recorded less than 10 hours.

Q-score calculation is implemented as a plugin to UCSF chimera, and is available from the following website: https://cryoem.slac.stanford.edu/ncmi/resources/software.

## Acknowledgements

This research has been supported by NIH grants (R01GM079429, P41GM103832, and S10OD021600). Molecular graphics and analyses were performed with the UCSF Chimera package. Chimera is developed by the Resource for Biocomputing, Visualization, and Informatics at the University of California, San Francisco (supported by NIGMS P41GM103311).

## Author contributions

G.P. conceived Q-scores, implemented the software and performed all the testing. W.C. came up with the term “Q-score”. K.Z. collected the images and reconstructed the maps (EMDB: 20026, 20027, 20028). Z.S. and S.L. provided additional data (not shown) for testing the Q-score. M.F.S. and W.C. contributed the discussion during the development. G.P. wrote the manuscript with inputs from other authors.

